# SHEPHARD: a modular and extensible software architecture for analyzing and annotating large protein datasets

**DOI:** 10.1101/2022.09.18.508433

**Authors:** Garrett M. Ginell, Aidan J. Flynn, Alex S. Holehouse

**Affiliations:** Department of Biochemistry and Molecular Biophysics, Washington University School of Medicine, St. Louis, MO, USA

## Abstract

The emergence of high-throughput experiments and high-resolution computational predictions has led to an explosion in the quality and volume of protein sequence annotations at proteomic scales. Unfortunately, integrating and analyzing complex sequence annotations remains logistically challenging. Here we present SHEPHARD, a software package that makes large-scale integrative protein bioinformatics trivial. SHEPHARD is provided as a stand-alone package and with a pre-compiled set of human annotations in a Google Colab notebook.

## INTRODUCTION

Over the last two decades, high-throughput experiments have enabled the acquisition of large datasets that offer insight into biologically important features for thousands of proteins simultaneously^1–4^. When combined with traditional and deep-learning-based computational approaches, proteome-wide annotation enables the generation of high-dimensional data ripe for further analysis ^5–9^ (**Fig. 1A**). If integrated, these datasets can be used to ask large-scale statistical questions on the relationship(s) between different types of annotations. These analyses can generate and test novel hypotheses, extracting additional value from previously published data in ways the original authors may have never anticipated.

**Figure 1.**
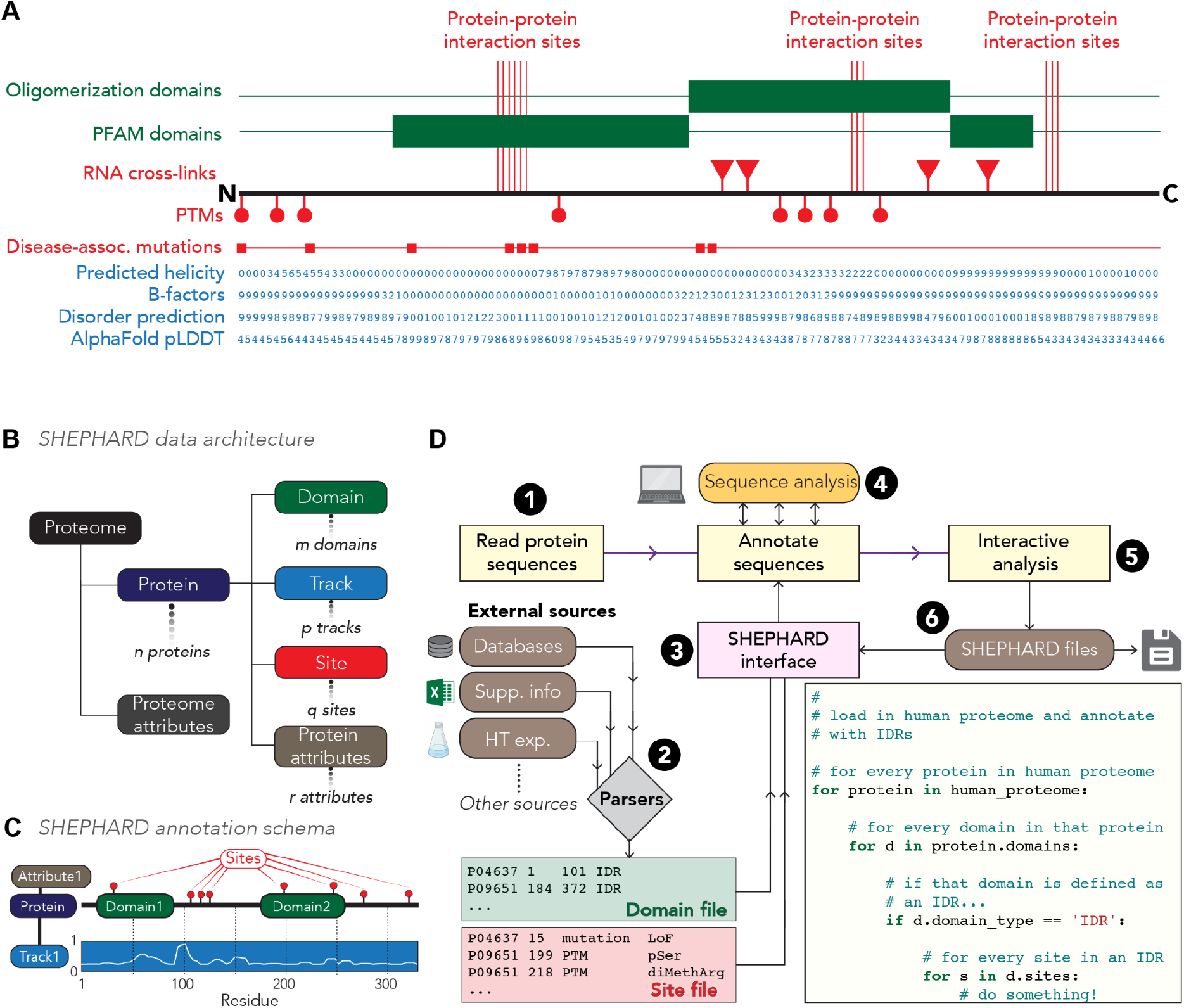
**(A)** Protein annotations span a range of flavors and types. **(B, C)** SHEPHARD uses four protein-level annotations, and multiple Proteins are organized in Proteomes. **(D)** SHEPHARD provides an ecosystem of tools for reading, analyzing, and writing annotations to facilitate syntactically simple analysis of large protein datasets.

Despite remarkable progress in data generation and acquisition, downstream processing and integration are often treated as an afterthought. Proteome-wide datasets that require incredible resources and effort to generate are often deposited in poorly-labeled Excel spreadsheets or hard-to-parse text files. In particular, the ability to cross-reference across many different types of annotations raises several practical challenges, including data cleaning, developing and applying appropriate data structures, and the portability and reliability of analysis code. These issues contribute to a scenario where large-scale analyses are often performed in a relatively *ad hoc* way. Researchers often invest substantial resources in building robust frameworks. Alternatively, they hack together bespoke, one-off pipelines, which may suffer from integrity, completeness, or reproducibility issues.

We have addressed this challenge by developing a modular and extensible software architecture for large-scale and high-throughput protein sequence analysis. SHEPHARD is a Python-based general-purpose hierarchical framework that facilitates reproducible, reliable, and high-throughput analysis of complex numerical and symbolic protein annotations at proteome-wide scales.

## RESULTS

SHEPHARD addresses three main challenges in the context of data integration and analysis. First, SHEPHARD enables a clear and syntactically simple programming interface for asking broad, integrative statistical questions across large datasets. Second, SHEPHARD makes it easy to read annotations from external sources and write annotations to files, making it easy to integrate into existing workflows. Third, we provide SHEPHARD as both a locally-installable Python package and Google Colab notebooks. These notebooks include various pre-loaded annotations for the human proteome. Taken together, SHEPHARD provides features for both seasoned bioinformaticians and novice users alike.

SHEPHARD stores data in an object-oriented hierarchical format where the base container is a Proteome. Proteomes contain one or more Proteins, and each Protein can be annotated with Domains, Sites, or Tracks (**Fig. 1B, C**). Proteomes and any associated annotations are read in and written out via a set of routines that provides an interface between the outside world and SHEPHARD. To ensure SHEPHARD-associated annotations are easy to generate, read, and interpret, we have introduced a simple tab-separated schema for defining SHEPHARD-associated data. Using a simple text-based format that can be read and written by commonly-used software (e.g., MS Excel) is a deliberate choice to ensure SHEPHARD-associated analyses remain accessible to anyone. Having introduced the conceptual and practical features that SHEPHARD addresses, the remainder of this report illustrates the types of integrative questions SHEPHARD makes easy to ask and answer across the human proteome.

Intrinsically disordered regions (IDRs) often contain post-translational modification (PTM) sites (**Table S1, S2**)^10^. We wondered if this observation reflects a *bona fide* preference of modifying enzymes for IDRs. Conversely, this result may be a convolution of sequence composition bias and differences in solvent accessibility for residues in IDRs vs. folded domains (**Fig. 2A**). To answer this, we combined proteome-wide per-residue binding accessibility data based on AlphaFold2 predicted structures with IDR and PTM annotations. These analyses reveal that after correcting for compositional biases and solvent accessibility, many (but not all) types of modifications remained enriched in IDRs, in line with recent work (**Fig. 2B**)^11^. For example, around 23% of serine residues in IDRs are phosphorylated compared to just 12% of solvent-accessible serine residues in folded domains. Moreover, most modifications occur in either disordered regions, flexible loops with no defined secondary structure, or alpha helices, suggesting these types of substrates are generally preferred (**Fig. S1, S2**).

**Figure 2.**
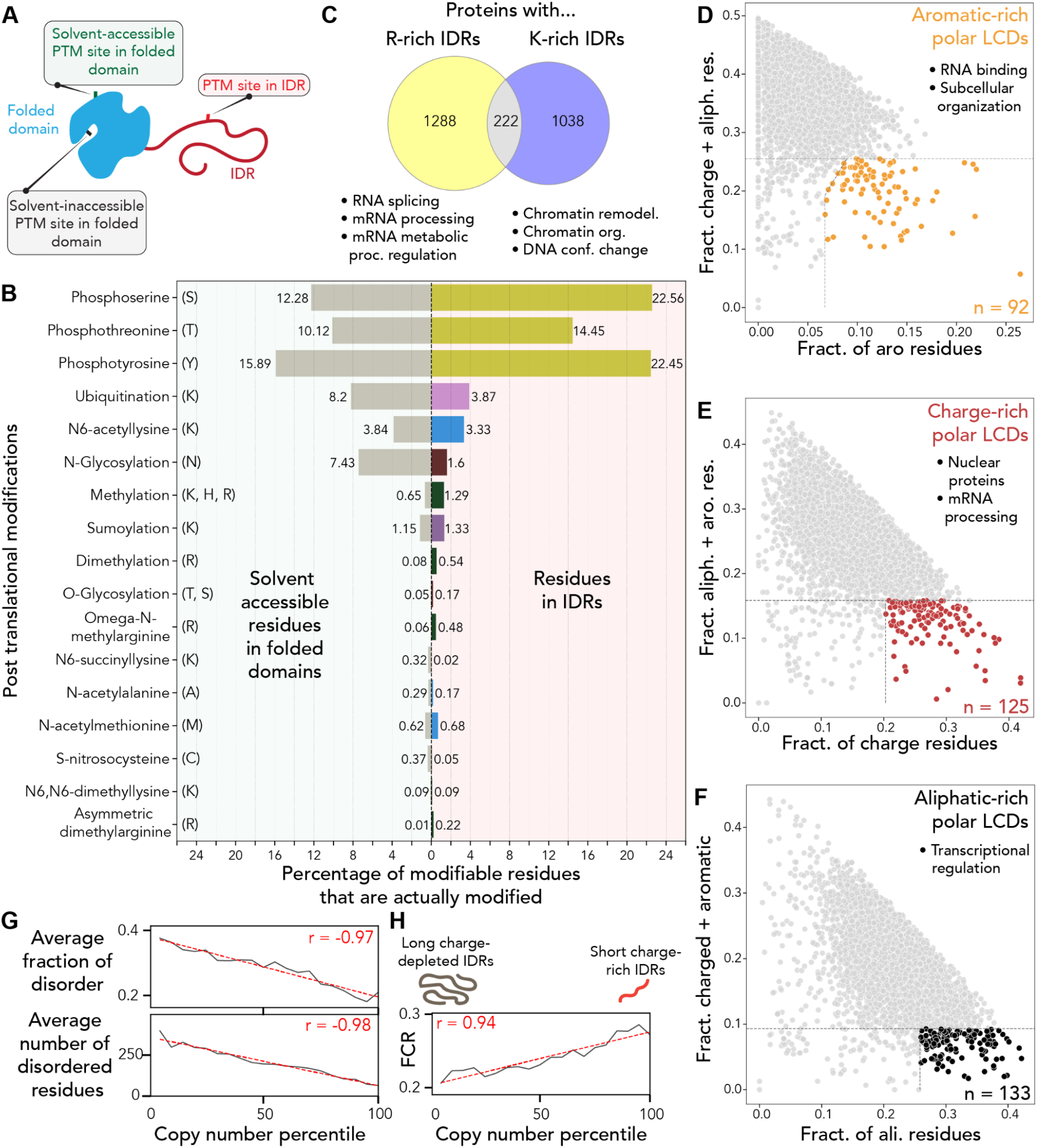
**(A)** Schematic of different types of PTM site types. **(B)** After accounting for structural context, residues in IDRs are still preferentially modified by PTMs. **(C)** Proteins with arginine or lysine-rich IDRs are largely non-overlapping in function and have distinct nucleic-acid binding preferences. **(D,E,F)** Proteins with polar-rich low-complexity domains enriched for specific types of chemistry have distinct functional enrichments. **(G)** Highly expressed human proteins tend to be depleted for IDRs (on average) (p < 0.0001). **(H)** IDRs in highly expressed human proteins tend to be more highly charged (p < 0.0001).

While IDRs are often treated as a monolithic class of domains, IDR conformational behavior and function can be influenced or even determined by amino acid sequence composition^12,13^. As such, we examined functional differences in IDRs enriched for either arginine or lysine, amino acids with different positively charged sidechains. While proteins with arginine-rich IDRs were enriched for RNA binding functions, those with lysine-rich IDRs were enriched for DNA binding (**Fig. 2C**, **Table S3, S4**). This analysis reveals interpretable chemical differences in nucleic-acid interaction preferences, mirroring previous demonstrations of a clear difference in higher-order assemblies driven by arginine or lysine-rich peptides^14^.

Motivated by our results from examining arginine and lysine-rich IDRs, we wondered if distinct flavors of polar-rich low complexity domains (pLCDs) might be associated with specific functions. pLCDs are regions in IDRs enriched in polar amino acids (glycine, serine, threonine, glutamine, asparagine, and here also proline)^15,16^. Given the polar nature of polypeptide backbones, pLCDs present a chemically homogenous scaffold that could be punctuated by other types of orthogonal chemistry. The kinds of sequence chemistry encoded by the remaining natural twenty amino acids can be broadly categorized as aliphatic, aromatic, or charged^17^. We identified pLCDs and filtered for those depleted in two of these three types of chemistry but enriched for the third. This revealed a clear difference in the types of proteins associated with chemically-distinct pLCDs (**Fig. 2D, E, F**). Aromatic-rich pLCDs (n=89) are enriched for RNA binding proteins as well as proteins involved in cellular structural components (intermediate filaments, nuclear pore complex, adhesion junction proteins) (**Table S5**). Charge-rich pLCDs (n=106) are largely enriched in nuclear proteins across a range of functions (notably RNA splicing and RNA processing) **Table S6**). Finally, aliphatic-rich pLCDs (n=117) are strongly enriched in proteins associated with transcriptional regulation (including transcription factors and chromatin binding proteins) (**Table S7**). Our results are consistent with a model whereby IDR chemical context can prime IDRs for specific functions.

Finally, prompted by prior work linking protein disorder to dosage-dependent toxicity, we wondered how the presence of disordered regions might correlate with protein abundance^18^. To our surprise, we observed a strong negative correlation between human protein copy number obtained by quantitative mass spectrometry and disorder (**Fig. 2G**). Encouragingly, this trend was reproduced across several other organisms with data from independent experiments (**Fig. S3, S4**). Intriguingly, across the human and yeast proteomes, we observed a strong correlation between how charged an IDR is and the copy number (**Fig. 2H**). IDRs in highly-abundant human proteins tend to be more highly charged. As such, we speculate that solubility – as determined by the fraction of charged residues – may play a role in defining the fitness advantage/defect associated with the presence of a given IDR. Our simple interpretation from these data is that long uncharged IDRs are generally more harmful than short, charged IDRs.

## CONCLUSION

SHEPHARD enables integrative proteome-wide bioinformatic analysis trivial for both local analysis pipelines and novice users via pre-defined Colab notebooks. To demonstrate SHEPHARD’s capacity, we have provided examples where annotations from AlphaFold2, integrative post-translational modification experiments, predicted disorder, sequence chemistry, and protein abundance data are seamlessly integrated. In summary, SHEPHARD offers a convenient resource for those working on understanding protein-function relationships at scale in a manner that is distributable, reproducible, and reliable.

## METHODS IN BRIEF

SHEPHARD is written in the Python programming language (https://www.python.org/) with limited core dependencies. Specifically, SHEPHARD requires Numpy^19,20^, a well-established library for numerical computing in Python, and protfasta, a high-performance FASTA parser developed explicitly for large protein datasets^21^.

SHEPHARD can be installed from the Python packaging index using the package installer for Python (**pip**) tool. Specifically, when run from the terminal, installation can be achieved using the following command:

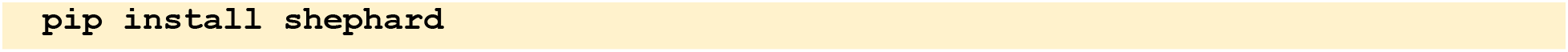

Documentation for using SHEPHARD is available at https://shephard.readthedocs.io/, with various examples and google-colab notebooks linked in the DATA AVAILABILITY section below.

Various annotations were used for the analysis performed here. These include proteome-wide sequence information provided from UniProt^8^, structural predictions from AlphaFold2^5,22^, post-translational modification data from ProteomeScout^23^, and gene ontology data from PANTHER^24^. In addition, protein abundance data were obtained for several different organisms, including humans^25^, *A. thaliana^26^, D. melongaster^27^, E. coli^28,29^, S. pombe^30^*, and *S. cerevisiae*^31,32^.

In addition to these sources of raw data, additional tools for inferring or interpreting sequence or structural data were also used. These include predicted disordered regions using metapredict (V2)^33^, prion-like domains identified using PLAAC^34^, low-complexity sequences identified using the approach developed by Gutierrez *et al.^15^*, and structural analysis using MDTraj^35^ and SOURSOP (https://soursop.readthedocs.io/). The structural classification was done using DSSP^36^ or DISICL^37,38^. Code for generating all figures and additional analysis are provided at the manuscript’s main GitHub repository (see **DATA AVAILABILITY**). A more extended methods section is provided in the supplementary information.

## ACKNOWLEDGEMENTS

This work was supported by Dewpoint Therapeutics, NSF Grant 2128068, and the Longer Life Foundation: an RGA/Washington University Collaboration. We thank all members of the Holehouse lab for comments and feedback and members of the Pappu lab for early discussion and thoughts on development.

## DATA AVAILABILITY

All data and code used to perform all analyses in this report can be found at https://github.com/holehouse-lab/supportingdata/tree/master/2022/ginell_2022. SHEPHARD documentation can be found at https://shephard.readthedocs.io/ and Colab notebooks can be found at https://github.com/holehouse-lab/shephard-colab. The SHEPHARD itself can be downloaded and installed from PyPI (https://pypi.org/project/shephard/) or from GitHub (https://github.com/holehouse-lab/shephard).

