## Supporting information for "SHEPHARD: a modular and extensible software architecture for analyzing and annotating large protein datasets"

### 1. SHEPHARD DESIGN DECISIONS

SHEPHARD is written in the Python programming language (https://www.python.org/) with limited core dependencies. Specifically, SHEPHARD requires Numpy[1,2], a well-established library for numerical computing in Python, and protfasta, a high-performance FASTA parser developed explicitly for large protein datasets[3]. Beyond these two external packages, our goal is to limit hard dependencies to core components of the Python scientific computing stack. The goal is to ensure the installation of SHEPHARD is as easy as possible.

SHEPHARD is built as a set of loosely coupled components. At its core, SHEPHARD contains a set of hierarchically-linked and self-referential data storage objects. These objects are **Proteomes**, **Proteins**, **Tracks**, **Domains**, and **Sites**. Each Proteome object stores zero or more Protein objects. Each Protein object stores zero or more Track objects, Domain objects, or Site objects. Tracks, Domains, and Sites are all aware of the Protein they originate from, such that analysis across all Sites, Tracks, or Domains can be performed as easily as it can be done over every Protein. In addition, Proteomes, Proteins, Sites, and Domains can have attributes associated with them. These attributes are essentially key-value pairs where the key defines an attribute name, and the value can be almost anything. Because arbitrary attributes can be annotated for all object types except Tracks, it is possible to extend annotations of a protein to whatever specific type of analysis is of interest. For example, SHEPHARD can be used to work with protein simulation data, where Protein attributes map to trajectories, or to protein cross-linking datasets, where site attributes include lists of cross-linked Sites (residues) from other proteins to construct interactome networks.

SHEPHARD data objects can be added and removed as analysis is performed. Tracks, Sites, and Domains can be added and removed from Proteins, and Proteins added or removed from Proteomes. This enables changes to data structures to be made during analysis, facilitating complex unidirectional pipelines that generate *de novo* annotations, prune datasets, or augment existing annotations.

A major advantage of casting protein annotations into one of four distinct types (Domains, Sites, Tracks, or Attributes) is that essentially identical code can often be re-used to analyze very different questions. By presenting a common style of interface for each of these objects, analogous design patterns can be learned for analyzing Sites and then applied to Domains (for example).

***Sequence indexing***

Python indexes from 0 and is exclusive (i.e., in so-called "slice syntax," the string `"abcde"` sliced as `abcde[1:2]` would return `"b"`). However, in biology (and in natural counting), we typically index from 1 and use an inclusive convention; that is, a domain that straddles residues 1-20 starts at the first residue (residue 1) and includes the $20^{th}$ (residue 20). SHEPHARD assumes this second indexing convention - that is, the first residue in a sequence is 1, and requesting subregions of sequences will honor this and include the residue associated with the end position. This decision avoids the need for users to perform offset corrections and makes it much easier to relate position in SHEPHARD to real-world residue numbering in sequence annotations. It also means if sequence subregions are to be excised, we provide a `get_sequence_region(start, end)` function in the Protein class.

***Interfaces***

In addition to the core data objects, SHEPHARD makes use of an **`interfaces`** module to read and write data to and from core data objects. Interfaces are a general design pattern from software engineering in which a common input format or structure is guaranteed to be readable by the associated software. In effect, an Interface provides a contract between the inner code of SHEPHARD and the outside world. This means that to load *any* type of data into SHEPHARD, the user need only transform their data into a datatype that is compliant with the interface. The alternative approach would be to build custom parsers for each data type, which places the burden of file parsing in the hands of the software developer and away from the expertise associated with the data. This is neither sustainable nor practical.

***Application programming interfaces (APIs)***

In addition to the ability to read/write SHEPHARD compliant annotations, SHEPHARD also provides an application programming interface (API) for interacting with specific external packages or types of data. The inclusion of an API module provides us with the ability to extend SHEPHARD to provide dedicated functionality for specific types of input, output, or analyses that go beyond the core data structures and interfaces. At the time of writing, the APIs include those for working with generic FASTA files and those for working with FASTA files obtained from UniProt. We anticipate adding additional APIs as the need arises.

***Sanity checking***

As a final note, SHEPHARD employs a 'safety first' approach to data parsing. That is, by default, operations that would overwrite data or lead to inconsistent behavior raise exceptions (errors). This behavior can be turned off on a case-by-case basis using the `safe` keyword. This design decision helps catch unexpected errors, conceptually flawed data types, or other problematic issues that may otherwise be silently missed.

### UNDERLYING SOFTWARE ARCHITECTURE

#### *Data types and data structure*

SHEPHARD implements a hierarchical data structure with datatypes that map to easily-interpretable protein features.

The base object in SHEPHARD is a ***Proteome object***. A Proteome is a collection of zero or more proteins. As well as providing a general organizing collection for Protein objects, Proteins enable easy access to a range of functionality. This includes the ability to retrieve all Domains or Sites of a certain type across all proteins, as well as retrieve the list of unique domain or site types found in the proteome. Proteome objects are also generators for proteins, meaning in Python syntax, one can iterate across the proteins in a Proteome simply by iterating over a for-loop.

***Protein objects*** possess information on an individual protein. This includes the underlying amino acid sequence, a unique identifier, a protein name, and zero or more annotations with Tracks, Domains, Sites, and Attributes. In addition, proteins contain a set of functions for excise sequence regions and contain the ability to add/remove Tracks, Domains, Sites, or Attributes.

***Track objects*** contain vectorial information that maps to each residue along a protein sequence and can encode either numerical (a values track) or symbolic information (a symbols track). A protein can have multiple tracks, but all tracks must match the sequence length. Tracks could be used to encode per-residue predictions, perform sequence coarse-graining, or encode experimental information in which a single value per-residue is generated (e.g., NMR chemical shifts, crystallographic B factors, CryoEM resolution, *etc.*)

***Domain objects*** describe discrete sub-regions within a protein and are defined by a start and end location, as well as a domain type. They are automatically named using a convention of `${domain_type}_${start}_${end}`. By default, two domains with identical types, start and end positions, are considered overlapping and will trigger an exception, but if the types are different, no such issue occurs. This is to avoid a scenario where multiple redundant domains are added, although if this is desired, this can be overridden using the `autoname` argument when adding domains. SHEPHARD also provides functionality to automatically construct domains based on Tracks, enabling the automatic discretization of continuous data into defined subregions. Domains are appropriate for structural information (e.g., annotated folded domains, helices, *etc.*), defined functional domains, or shorter sequence motifs.

***Site objects*** map to a single position within a protein sequence. A Site is defined by a position, a site type, the associated protein, and symbolic or numerical data. Multiple sites can coexist at the same residue. Sites are appropriate for specific positional information such as single point mutations, post-translational modifications, or known binding residues.

For Domains, information from Tracks or Sites found within the domain can be directly extracted. Similarly, for Sites, Track-specific data can be obtained, as can Domains that encompass a Site. We can interrogate both Proteomes and Proteins for Domains or Sites of a specific type, and we can identify proteins from which Sites of a specific type originate. In effect, the SHEPHARD data structure offers a relational database grounded using a Primary Key (`protein.unique_ID`) associated with each protein, yet offering syntactically intuitive ways to query and analyze in what feels like conventional Python code.

***Generating SHEPHARD Proteomes***

Generating and populating proteomes into SHEPHARD can be achieved in several ways.

The recommended route is to start with a FASTA file with one or more sequence records (**Fig. S6**). The FASTA format stipulates that each sequence is demarked with a header line that starts with a caret (>) character followed by identifying information. FASTA files can be used to create new Proteomes using the FASTA API (`shephard.apis.fasta`). In addition, if the FASTA file was obtained from UniProt (http://uniprot.org/), then the UniProt API (`shephard.apis.uniprot`) can be used to parse a FASTA file such that UniProt accession is used as the protein unique ID and the protein name is parsed from the FASTA header.

In addition to loading data from FASTA files, SHEPHARD enables previously-generated SHEPHARD protein files to be read using the proteins SHEPHARD interface (`shephard.interfaces.si_proteins`). SHEPHARD-formatted protein files include a single line per protein, as described in the reference implementation for the proteins files (defined in the online documentation https://shephard.readthedocs.io/en/latest/shephard_file_types.html). As with FASTA files, these files can be read to generate a new Proteome object in a single line of code.

Finally, proteomes can be constructed programmatically. A new Proteome object can be created using an empty constructor, and then proteins added using the (`add_protein()`) function. New proteins require a sequence, a unique ID, and a name and can have zero or more attributes.

***Importing SHEPHARD protein annotations***

Once a Proteome has been generated, it can be annotated with Tracks, Domains, Sites, or Attributes. In all four cases, these annotations can be done using functions from the appropriate interface model in (`shephard.interfaces`). Interfaces implement consistent, stateless functions that enable the reading (and writing) of SHEPHARD data from (and to) a standardized file format. This file format SHEPHARD uses by default is a simple tab-separated text format that is easily generated or readable using commonly used software (e.g., Excel). The objective is to make it easy to read (and write) data into (and out of) SHEPHARD.

By way of example, for annotating a Proteome object with domains, one would use the `shephard.interfaces.si_domains.add_domains_from_file()` function. This function takes two arguments: a Proteome object and the path to the Domains file. During parsing of the Domains file, SHEPHARD ensures that annotations do not extend beyond the positions available in a domain, and only annotations for Protein objects found in the Proteome are added.

In addition to reading annotations from an annotation file, `shephard.interfaces` enable dictionaries of annotations to be used to annotate proteomes programmatically. The specifics for dictionary-derived annotations are available in the SHEPHARD reference documentation. The main advantage here is that using a dictionary-based annotation enables dynamic annotations to be added, allowing SHEPHARD to be integrated into an existing software pipeline without the need for writing to disk.

***Exporting SHEPHARD protein sequence information***
As with reading protein sequence information, writing sequence information can be achieved in two different ways. Proteome objects can be exported to FASTA file via the FASTA ASI (`api.fasta`).

***Exporting SHEPHARD annotations***
Annotations from a Proteome object can be exported for future use or distribution. There are two ways this can be done. In one approach, the Proteome object is passed into one of the `write_<annotation>` functions (e.g., `shephard.interfaces.si_sites.write_sites()`). These functions take a Proteome object, an output path, and then enable specific types of annotations to be written based on (for example) Site type or Domain Type.

The alternative approach is to manually generate a Python list of the annotations of interest and then pass this list of annotation objects to a `write_<x>_from_list` function. For example, to write a list of sites the function `shephard.interfaces.si_sites.write_sites_from_list()` may be used. This approach provides much finer-grain control over which annotations are actually written - for example, one could iterate through all the sites in a Proteome and add only those from proteins under some threshold number of residues to a new list (using standard Python list syntax of `append()`).

These two modes of data export enable the two standard scenarios: either a user wants to export all the annotations of a specific type or class to facilitate full reproducibility, or a user wants to export a very specific set of annotations for further analysis downstream.

***SHEPHARD Application Programming Interfaces (APIs)***
While SHEPHARD guarantees the ability to read and write in its standardized formats, some additional formats are sufficiently broadly used that they merit their own specific APIs for reading and writing. As introduced above, FASTA files are the *de facto* standard format for sharing protein sequence information, and, as such, ensuring SHEPHARD provides robust capabilities for reading and writing FASTA files is important.

For standard FASTA file reading/writing, the functions encapsulated by the fasta module should be sufficient (e.g., `shephard.apis.fasta`). In addition, given we generally recommend working with data obtained from UniProt for proteome-wide analysis, we provided an UniProt API for reading/writing FASTA data obtained from UniProt (e.g., `shephard.apis.uniprot`). More broadly, we plan to add additional APIs for specific tools or packages as soft dependencies moving forward; i.e. APIs provide a means for SHEPHARD to include modular gateways to additional functionality. Moreover, APIs are not a core component of SHEPHARD and instead should be thought of as a convenient layer of regulation and functionality.

***SHEPHARD tools***

In addition to core functionality encoded within different SHEPHARD objects, we provide a set of stand-alone tools for manipulating SHEPHARD data. These modules include `shephard.tools.domain_tools`, `shephard.tools.sequence_tools`, `shephard.tools.site_tools` and `shephard.tools.track_tools`.

The tools modules share the fact that all functions are stateless, stand-alone, and do not modify the objects passed into them. In this way, tool functions enable the manipulation or analysis of existing data. SHEPHARD annotation objects make use of functions encoded in the tools modules, but in addition, certain functionality that may of use or of interest for those using SHEPHARD are encoded in the tools modules

In particular, the `shephard.tools.track_tools.build_track_from_domains()` function enables a set of domains to be converted into a track. The corollary is also possible with the `shephard.tools.domain_tools.build_domains_from_track()` function. This is particularly useful for identifying regions that can discretize a continuous (track) annotation into a binary (domain) annotation. For example, one could use this to identify 'domains' with a high propensity for some predictable value.

While the tools modules contain relatively few functions at the time of writing, our goal is to have these modules as a public-facing location for additional functions that may emerge. In this way, pull requests for new types of features can be implemented here, in a way that, by definition, does not impact core components of the SHEPHARD architecture. In much the same way that the APIs are designed to be modular and extensible, so too are the tools modules.

### 2. EXTENDED DISCUSSION OF PERFORMING ANALYSIS IN SHEPHARD

The power of SHEPHARD is in the ability to – in just a few lines of code – iterate over entire proteomes and analyze large-scale sequence annotations using the conventional syntax of Python programming. Rather than requiring users to become familiar with an entirely new interface, SHEPHARD is designed to operate in a manner that is consistent, robust, and easy to use. As well as making analysis straightforward, our hope is that this will allow users to write that is easy for non-experts to read and understand. Python benefits from being almost pseudocode-like at times, and the recommended iterative style of analysis in SHEPHARD is conducive to writing code that is easy for humans to read.

Upon reading sequence information into a Proteome, Proteins can be called or iterated over in a fashion analogous to a Python dictionary. By ensuring different annotations are aware of their protein of origin, complex, referential analysis pipelines can be constructed in a few lines of code. Moreover, we can examine the intersection between annotations by (for example) identifying Sites that fall within a specific type of domain or by building a Track object for a given protein based on the Protein's amino acid sequence by calling the `build_track()` function from the Protein of interest.

SHEPHARD's hierarchical framework also makes analyses that may previously have been cumbersome or difficult efficient and fast, both in execution and in ease of development. In particular, the ability to inherent cross-reference between different annotation types is a major advantage. For example, from a given domain, one can extract the underlying sequence (`domain.sequence`*)*, and sites found within that domain (`domain.sites`), or the track values or symbols associated with a domina are obtained using `domain.get_track_values()` or `domain.get_track_symbols()` functions. Several examples of analysis that make use of these features are illustrated in notebooks found at https://github.com/holehouse-lab/shephard-colab.

In addition to making use of built-in analysis functions, SHEPHARD enables Proteins, Proteomes, Tracks, Domains, and Sites can be annotated with attributes. Attributes are arbitrary key-value pairs that enable both data and functionality to be associated with an object. In particular, functions in Python are first-class objects. With that in mind, data objects can be annotated with novel analysis functions. This means custom analysis can be developed, annotated to a data object, and then performed *in situ* within an analysis pipeline. Various examples of SHEPHARD-based analysis are provided as both example notebooks, and in the code for this manuscript.

### 3. GOOGLE COLAB NOTEBOOKS

*Demo analysis*

In addition to a collection of example notebooks for SHEPHARD, we provide a set of Google-colab notebooks which are linked from https://github.com/holehouse-lab/shephard-colab. These notebooks are provided to illustrate interactive examples of the types of analysis SHEPHARD can provide.

*Human proteome notebook*

In addition to notebooks with simple examples, we also provide a notebook in which the human proteome is fully annotated. The annotations include post-translational modifications, intrinsically disordered regions, prion-like domains, per-residue secondary structure annotation, per-residue solvent accessibility scores, and protein copy number.

### 4. DATA

All the data and associated analysis scripts described below are provided in the main GitHub repository at

https://github.com/holehouse-lab/supportingdata/tree/master/2022/ginell_2022

#### *Human proteome*

The human proteome was obtained from UniProt in 2020 and used as a reference for this study [4]. In part, this helps maintain consistency with the AlphaFold2 data, which was obtained in July 2022 from the initial (V1) AlphaFold/EBI data deposition [5,6]. Note that we compared the predicted structures for the human proteome with the second AlphaFold2/EBI data deposition (December 2021) and found almost no changes - a full description of this is available under the `/misc` directory on the SHEPHARD manuscript github page

To ensure compatibility with the provided AlphaFold2 data, any proteins without an exact sequence match between the AlphaFold2 data and the model human proteome were filtered out. This left 20,061 proteins (98.4% of the original human proteome). This ultimately left 326,924 PTM annotations (99.8% of the original annotated PTMs). It is worth noting that the filtering here has no material impact on the results.

All Proteomes were similarly obtained from the UniProt Proteomes interface found at https://www.uniprot.org/proteomes.

Proteins with invalid amino acids were removed using protfasta [3] using the pfasta command-line tool. As an example:

```
pfasta --invalid-sequence remove raw_human_proteome.fasta -o human_proteome_validated.fasta
```

#### *Intrinsically disordered domains*

Intrinsically disordered regions were predicted using metapredict V2 [7,8].

Specifically, we used the command-line tool **`metapredict-predict-idrs`** with the **`--mode`** set to **`shephard-domains`** flag.

```
metapredict-predict-idrs human_proteome_validated.fasta --mode shephard-domains-uniprot \\
                         -o shprd_domains_metapredictv2.tsv --verbose
```

We note that proteome-wide disorder predictions can also be made using the metapredict Google colab notebook linked from https://metapredict.net/.

#### *Polar-rich low complexity domains (pLCDs)*

Polar-rich low complexity domains (pLCDs) were identified using the method developed by Gutierrez *et al.* [9]. Specifically, this approach identifies contiguous regions enriched in specific residues, enabling gap sizes and minimum region sizes to be defined. We selected regions in the human proteome rich in polar and proline residues (Q, S, G, N, T, & P), with a maximum interruption of 5 residues, a minimum domain size of 50 residues, and a fractional threshold of 0.5 or higher. Code for computing this is provided at the GitHub repository under `/misc/find_plcds/`, and the specific implementation is provided in the sequence analysis package, sparrow (https://github.com/idptools/sparrow).

For pLCDs, we then selected only LCDs that overlap with IDRs, and calculated the fractions of polar, aromatic, aliphatic, and charge residues in each region. Chemically distinct polar LCDs are then defined as LCD that are depleted in two of the chemical types but above a specific relative threshold in the chemical type of interest. For example, aromatic-rich pLCDs are depleted in charged and aliphatic residues and have at least 50% more aromatics relative to the fraction of charge and aliphatic residues combined.

In the spirit of completeness, we also calculated a large collection of chemically distinct LCDs, all of which are provided as a SHEPHARD domains file under `/misc/find_plcds/shprd_domains_human_LCDs_all.tsv`.

**Post-translational modifications (PTM)**
PTM data were obtained from ProteomeScout (downloaded summer 2022) and parsed using the ProteomeScoutAPI [10,11]. Specifically, PTM data were downloaded from https://proteomescout.wustl.edu/compendia.

**Structural annotations**
Structural annotations across the human proteome were based on predicted protein structures from AlphaFold2 [5,6]. The decision to use computed structural models for proteome-wide structural analysis reflects the convenience this affords with respect to guaranteed coverage, 1:1 mapping between sequence and structure, and the remarkable accuracy that AlphaFold2 offers. Classification into distinct structural subclasses was performed using the DISICL classification scheme developed by Nagy *et al.* and implemented in the SESCA package [12,13]. We also employed the DSSP algorithm to annotate the human proteome based on the AlphaFold2 annotations [14].

**Accessible residues**
Residue accessibility was calculated using SOURSOP (https://soursop.readthedocs.io/) and MDTraj [15], using a solvent probe radius of 7 Å. Code for computing this is provided at the GitHub repository under `/PTM_af2_analysis/residue_accessibility/`.

**Gene ontology enrichment**
Gene ontology enrichment was performed using PANTHER [16]. In all cases, the complete set of UniProt IDs associated with compositional filtering was uploaded, and enrichment against the human proteome was calculated.

**Protein abundance data**

Protein abundance data was included for humans [17], *A. thaliana* [18], *D. melongaster* [19], *E. coli* [20,21], *S. pombe* [22], and *S. cerevisiae* [23,24]. To calculate correlations, we took the set of proteins for which abundance data were available and separated them into ventiles (twenty equally-spaced groups, rank-ordered from lowest to highest expression levels). The use of ventiles was primarily to ensure we also had groups with sufficiently large numbers of proteins, although where accessible changing the number of bins did not alter the results. For each ventile, we calculated the average fraction disorder of proteins, the average number of disordered residues, and the average fraction of charged residues (FCR) for disordered regions. The correlation between ventiles and the calculated property was then computed using the Pearson's correlation coefficient. Code for computing this is provided in the GitHub repository under `IDR_copy_number/copy_number_analysis.ipynb`.

**Prion-like domains**

Prion-like domains were predicted using the PLAAC algorithm [25,26]. The PLAAC source code was obtained from https://github.com/whitehead/plaac, and run using default parameters to analyze the human proteome.

**Extended Acknowledgements**

We thank the members of both the Holehouse lab and Pappu Lab for ongoing helpful discussions and feedback over the course of the development of the package. In particular, we thank Jared M. Lalmansingh for their input during the package's initial development. We also thank D. Allan Drummond and Stephen Fried for helpful discussions. A special thanks to Shubhanjali Minhas for designing the SHEPHARD logo.

### SUPPLEMENTARY TABLES

| | Number of residues | Number of PTM sites | Percentage of residues that are modified |
|---|---|---|---|
| Human proteome | 10,483,347 | 312,745 | 3.0 % |
| Human IDRs | 3,439,190 | 154,758 | 4.5 % |

**Table S1.** IDRs are disproportionately post-translationally modified. On average, 4.5% of all disordered residues have one or more post-translational modifications. Moreover, 50% of all residues that have one or more modifications are found in IDRs, despite the fact that 33% of residues in the human proteome are found in IDRs. Note that PTM sites are defined as residues with one or more post-translational modification; i.e. a residue that received two different types of modifications would be counted once.

| Post-translational modification | Count |
|---|---|
| Phosphoserine | 131,681 |
| Phosphothreonine | 54,168 |
| Phosphotyrosine | 38,412 |
| Ubiquitination | 38,196 |
| N6-acetyllysine | 20,331 |
| N-Glycosylation | 13,882 |
| Methylation | 11,535 |
| Sumoylation | 6,528 |
| Dimethylation | 1,375 |
| O-Glycosylation | 1,255 |
| Omega-N-methylarginine | 1,154 |
| N6-succinyllysine | 1,152 |
| N-acetylalanine | 949 |
| N-acetylmethionine | 820 |
| S-nitrosocysteine | 729 |
| N6,N6-dimethyllysine | 504 |
| Asymmetric dimethylarginine | 477 |

**Table S2.** PTMs of interest across the human proteome. We focus on the most numerous 17 types of modifications.

| Term | Total | Actual | Expected | Enrich. | Raw P value | FDR |
|---|---|---|---|---|---|---|
| regulation of RNA splicing (GO:0043484) | 75 | 37 | 4.28 | 8.64 | 8.47E-20 | 2.35E-17 |
| mRNA splicing, via spliceosome (GO:0000398) | 176 | 66 | 10.05 | 6.57 | 9.32E-29 | 5.18E-26 |
| RNA splicing, via transesterification reactions with bulged adenosine as nucleophile (GO:0000377) | 176 | 66 | 10.05 | 6.57 | 9.32E-29 | 4.14E-26 |
| RNA splicing, via transesterification reactions (GO:0000375) | 176 | 66 | 10.05 | 6.57 | 9.32E-29 | 3.45E-26 |
| RNA splicing (GO:0008380) | 221 | 77 | 12.62 | 6.1 | 1.23E-31 | 1.37E-28 |
| mRNA processing (GO:0006397) | 229 | 79 | 13.08 | 6.04 | 3.41E-32 | 7.58E-29 |
| regulation of mRNA metabolic process (GO:1903311) | 110 | 32 | 6.28 | 5.09 | 3.78E-12 | 5.25E-10 |
| mRNA metabolic process (GO:0016071) | 316 | 88 | 18.05 | 4.88 | 5.18E-30 | 3.84E-27 |
| calcium ion transmembrane transport (GO:0070588) | 80 | 19 | 4.57 | 4.16 | 1.25E-06 | 6.93E-05 |

**Table S3. GO enrichment terms for proteins with arginine-rich IDRs.** Proteins with arginine-rich IDRs are enriched for RNA-associated annotations, along with the curious inclusion of calcium ion transmembrane transport.

| Term | Total | Actual | Expected | Enrich. | Raw P value | FDR |
|---|---|---|---|---|---|---|
| chromatin remodeling (GO:0006338) | 77 | 30 | 3.84 | 7.81 | 1.26E-15 | 2.00E-13 |
| chromatin organization (GO:0006325) | 118 | 35 | 5.89 | 5.95 | 5.24E-15 | 7.75E-13 |
| DNA conformation change (GO:0071103) | 74 | 20 | 3.69 | 5.42 | 1.36E-08 | 5.30E-07 |
| DNA recombination (GO:0006310) | 115 | 26 | 5.74 | 4.53 | 2.48E-09 | 1.06E-07 |
| protein-DNA complex subunit organization (GO:0071824) | 83 | 18 | 4.14 | 4.35 | 1.17E-06 | 3.16E-05 |
| chromosome organization (GO:0051276) | 256 | 53 | 12.77 | 4.15 | 3.36E-16 | 5.73E-14 |

**Table S4. GO enrichment terms for proteins with lysine-rich IDRs**. Proteins with lysine-rich IDRs are enriched for DNA-associated annotations (notably those associated with chromatin and chromosomal annotations).

| | Proteins with aromatic-rich pLCDs | | | | | | |
|---|---|---|---|---|---|---|---|
| | **Term** | **Total** | **Actual** | **Expected** | **Enrich.** | **Raw P value** | **FDR** |
| **Cellular Component** | keratin filament (GO:0045095) | 31 | 4 | 0.13 | 30.54 | 1.39E-05 | 3.54E-03 |
| | nuclear pore (GO:0005643) | 47 | 4 | 0.2 | 20.14 | 6.28E-05 | 1.07E-02 |
| | intermediate filament (GO:0005882) | 50 | 4 | 0.21 | 18.93 | 7.88E-05 | 1.01E-02 |
| | intermediate filament cytoskeleton (GO:0045111) | 51 | 4 | 0.22 | 18.56 | 8.47E-05 | 8.65E-03 |
| **Molecular Function** | structural constituent of nuclear pore (GO:0017056) | 19 | 4 | 0.08 | 49.82 | 2.44E-06 | 4.43E-04 |
| | single-stranded RNA binding (GO:0003727) | 33 | 3 | 0.14 | 21.51 | 4.65E-04 | 3.62E-02 |
| | signal sequence binding (GO:0005048) | 33 | 3 | 0.14 | 21.51 | 4.65E-04 | 3.17E-02 |
| | RNA binding (GO:0003723) | 617 | 20 | 2.61 | 7.67 | 1.52E-12 | 8.33E-10 |
| | structural molecule activity (GO:0005198) | 230 | 7 | 0.97 | 7.2 | 6.38E-05 | 5.81E-03 |

**Table S5. GO enrichment terms for proteins with aromatic-rich polar-rich low-complexity domains (aromatic-rich pLCDs)**. Proteins identified include those associated with RNA binding, as well as those with structural roles in subcellular organization.

| | Proteins with charge-rich pLCDs | | | | | | |
|---|---|---|---|---|---|---|---|
| | **Term** | **Total** | **Actual** | **Expected** | **Enrich.** | **Raw P value** | **FDR** |
| **Biological process** | mRNA splicing, via spliceosome (GO:0000398) | 176 | 9 | 0.9 | 10.03 | 4.31E-07 | 2.39E-04 |
| | RNA splicing, via transesterification reactions (GO:0000375) | 176 | 9 | 0.9 | 10.03 | 4.31E-07 | 1.60E-04 |
| | mRNA processing (GO:0006397) | 229 | 11 | 1.17 | 9.42 | 3.88E-08 | 8.61E-05 |
| | RNA splicing (GO:0008380) | 221 | 10 | 1.13 | 8.87 | 2.83E-07 | 2.09E-04 |
| | mRNA metabolic process (GO:0016071) | 316 | 11 | 1.61 | 6.83 | 8.62E-07 | 2.73E-04 |
| | RNA processing (GO:0006396) | 481 | 12 | 2.45 | 4.89 | 7.67E-06 | 1.55E-03 |
| **Cellular component** | transcription elongation factor complex (GO:0008023) | 34 | 5 | 0.17 | 28.84 | 1.53E-06 | 1.31E-04 |
| | nuclear body (GO:0016604) | 50 | 6 | 0.25 | 23.53 | 3.90E-07 | 6.65E-05 |
| | nucleoplasm (GO:0005654) | 422 | 15 | 2.15 | 6.97 | 5.73E-09 | 1.46E-06 |
| | nuclear lumen (GO:0031981) | 632 | 15 | 3.22 | 4.65 | 9.35E-07 | 9.55E-05 |
| | membrane-enclosed lumen (GO:0031974) | 731 | 15 | 3.73 | 4.02 | 5.35E-06 | 3.91E-04 |
| | intracellular organelle lumen (GO:0070013) | 731 | 15 | 3.73 | 4.02 | 5.35E-06 | 3.42E-04 |
| | organelle lumen (GO:0043233) | 731 | 15 | 3.73 | 4.02 | 5.35E-06 | 3.04E-04 |

| | | | | | | | |
|---|---|---|---|---|---|---|---|
| **Molecular function** | transcription coregulator activity (GO:0003712) | 210 | 7 | 1.07 | 6.54 | 1.21E-04 | 2.20E-02 |
| | RNA binding (GO:0003723) | 617 | 16 | 3.15 | 5.08 | 1.22E-07 | 6.66E-05 |

**Table S6. GO enrichment terms for proteins with charge-rich polar-rich low-complexity domains (charge-rich pLCDs)**. Proteins identified include many associated with nuclear function and nuclear subcellular localization. These include RNA processing, splicing, and proteins associated with the nucleoplasm.

| | Proteins with aliphatic-rich pLCDs | | | | | | |
|---|---|---|---|---|---|---|---|
| | **Term** | **Total** | **Actual** | **Expected** | **Enrich.** | **Raw P value** | **FDR** |
| **Biological Process** | transcription by RNA polymerase II (GO:0006366) | 1438 | 51 | 8.17 | 6.24 | 9.80E-28 | 5.44E-25 |
| | regulation of transcription by RNA polymerase II (GO:0006357) | 1383 | 47 | 7.86 | 5.98 | 1.52E-24 | 1.77E-22 |
| | nucleic acid-templated transcription (GO:0097659) | 1790 | 58 | 10.17 | 5.7 | 3.79E-30 | 8.42E-27 |
| | RNA biosynthetic process (GO:0032774) | 1798 | 58 | 10.22 | 5.68 | 4.78E-30 | 3.53E-27 |
| | regulation of transcription, DNA-templated (GO:0006355) | 1713 | 53 | 9.73 | 5.44 | 3.37E-26 | 6.23E-24 |
| | regulation of RNA metabolic process (GO:0051252) | 1853 | 56 | 10.53 | 5.32 | 1.98E-27 | 8.81E-25 |
| | regulation of nucleobase-containing compound metabolic process (GO:0019219) | 1911 | 56 | 10.86 | 5.16 | 9.03E-27 | 2.23E-24 |
| | regulation of cellular macromolecule biosynthetic process (GO:2000112) | 1849 | 53 | 10.51 | 5.04 | 1.16E-24 | 1.61E-22 |
| | nucleobase-containing compound biosynthetic process (GO:0034654) | 2026 | 58 | 11.51 | 5.04 | 2.16E-27 | 8.00E-25 |
| | regulation of gene expression (GO:0010468) | 2106 | 57 | 11.97 | 4.76 | 1.28E-25 | 1.90E-23 |

| | | | | | | | |
|---|---|---|---|---|---|---|---|
| | RNA metabolic process (GO:0016070) | 2410 | 61 | 13.7 | 4.45 | 2.96E-26 | 5.97E-24 |
| **Molecular function** | transcription regulator activity (GO:0140110) | 1265 | 39 | 7.19 | 5.43 | 1.22E-18 | 3.32E-16 |
| | RNA polymerase II transcription regulatory region sequence-specific DNA binding (GO:0000977) | 1059 | 32 | 6.02 | 5.32 | 5.69E-15 | 4.44E-13 |
| | DNA binding (GO:0003677) | 1361 | 41 | 7.73 | 5.3 | 2.65E-19 | 1.44E-16 |
| | nucleic acid binding (GO:0003676) | 1974 | 46 | 11.22 | 4.1 | 1.60E-17 | 2.92E-15 |

**Table S7. GO enrichment terms for proteins with aliphatic-rich polar-rich low-complexity domains (aliphatic-rich pLCDs)**. Proteins identified include mostly those involved in transcriptional regulation across a range different processes, including RNA synthesis,

### SUPPLEMENTARY FIGURES

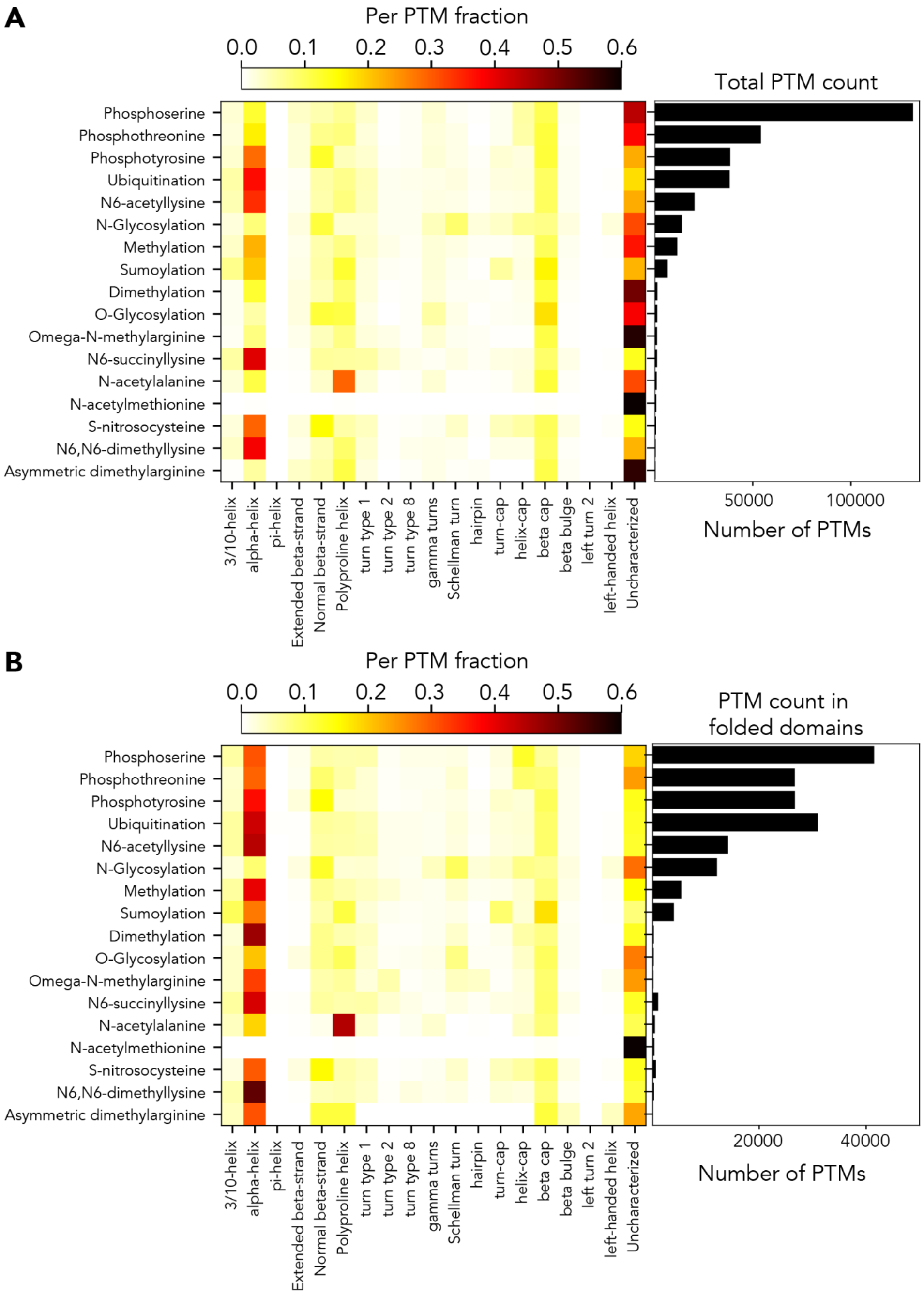


**Figure S1. PTMs show strong structural biases for the local structural context. (A)** PTMs overwhelmingly fall in regions that lack a defined secondary structure or alpha helices. **(B)** If only PTMs in folded regions are analyzed, similar trends are recovered, with the balance between short disordered loops and alpha helices largely swapping.

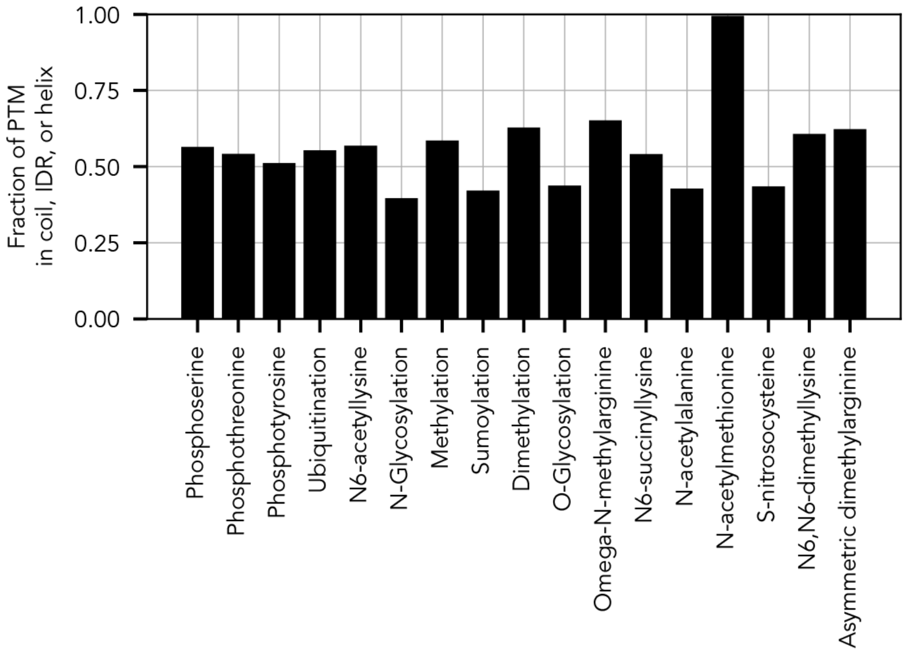


**Figure S2. Fraction of PTMs in solvent-accessible structural contexts.** For each type of modification, we determined what fraction of that residue was found in one of (1) unstructured/unclassified regions, (2) disordered regions, or (3) helices. For most PTMs, the majority were found in one of these three structural contexts.

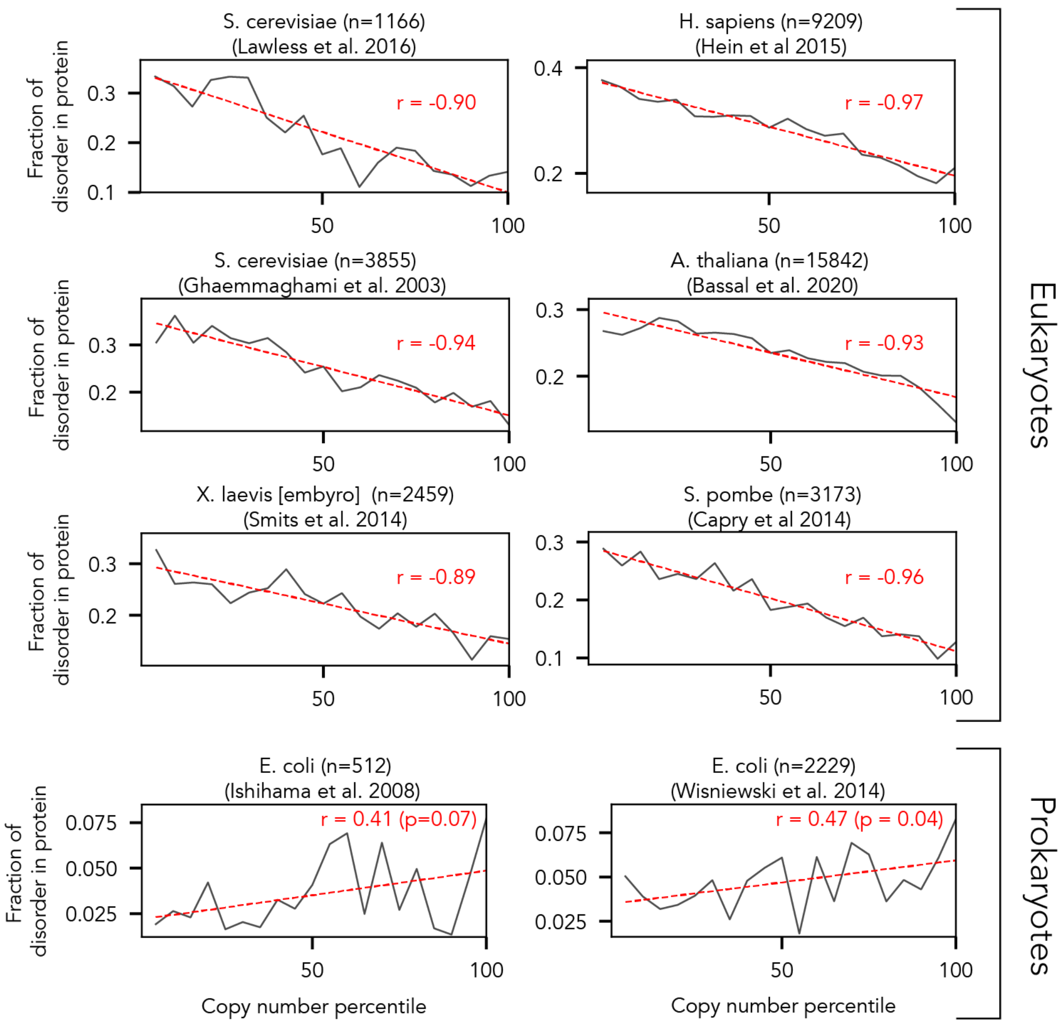


**Figure S3. Copy number vs. the average fraction of disorder across eight different mass spectrometry datasets**. Taking supplementary information from eight different studies, we correlated the average fraction of protein disorder in proteins within each ventile against the copy number ventiles (represented as percentiles for convenience), with correlations calculated using Pearson's correlation coefficient. While the eukaryotes examined consistently show a negative correlation between copy number and fraction of disorder, prokaryotes show no statistically significant trends ($p > 0.01$), a result that likely reflects the comparably lower propensity of disorder in prokaryotes.

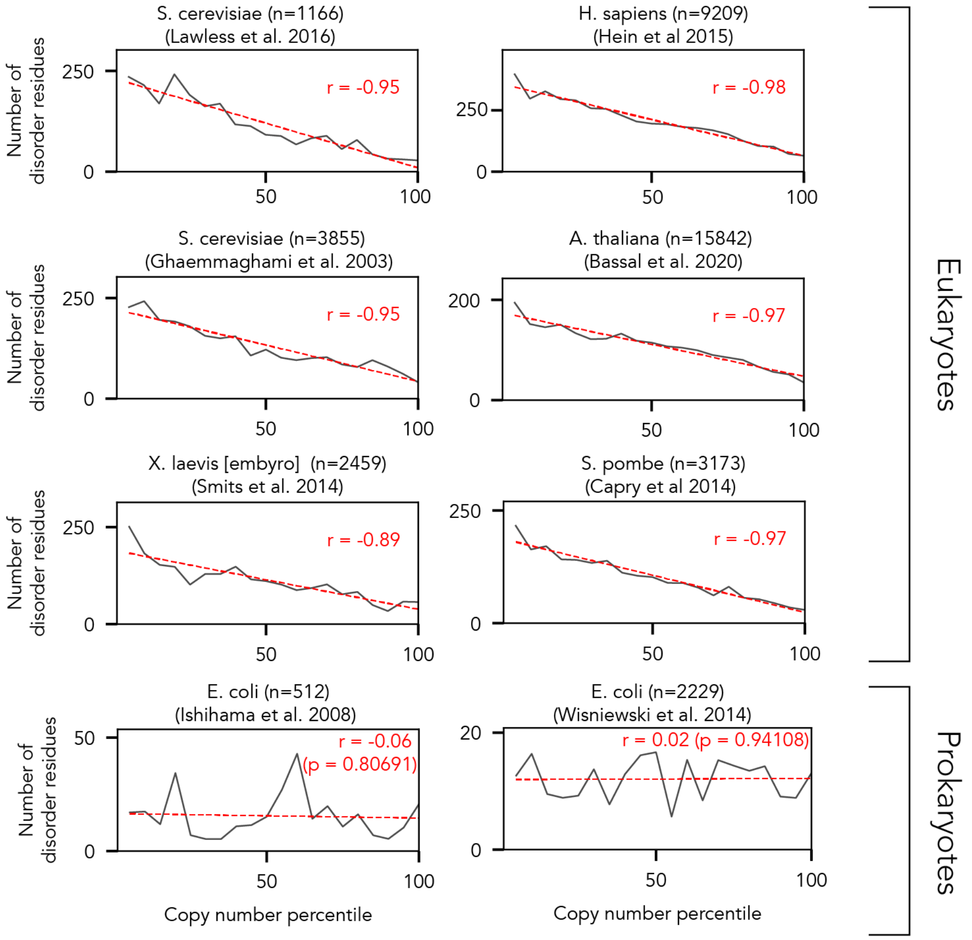


**Figure S4. Copy number vs. the average number of disorder residues across eight different mass spectrometry datasets**. Taking supplementary information from eight different studies, we correlated the average number of disorder residues in proteins within each ventile against the copy number ventiles (represented as percentiles for convenience), with correlations calculated using Pearson's correlation coefficient. While the eukaryotes examined consistently show a negative correlation between copy number and number of disordered residues, prokaryotes show no statistically significant trends ($p > 0.01$), a result that likely reflects the comparably lower propensity of disorder in prokaryotes.

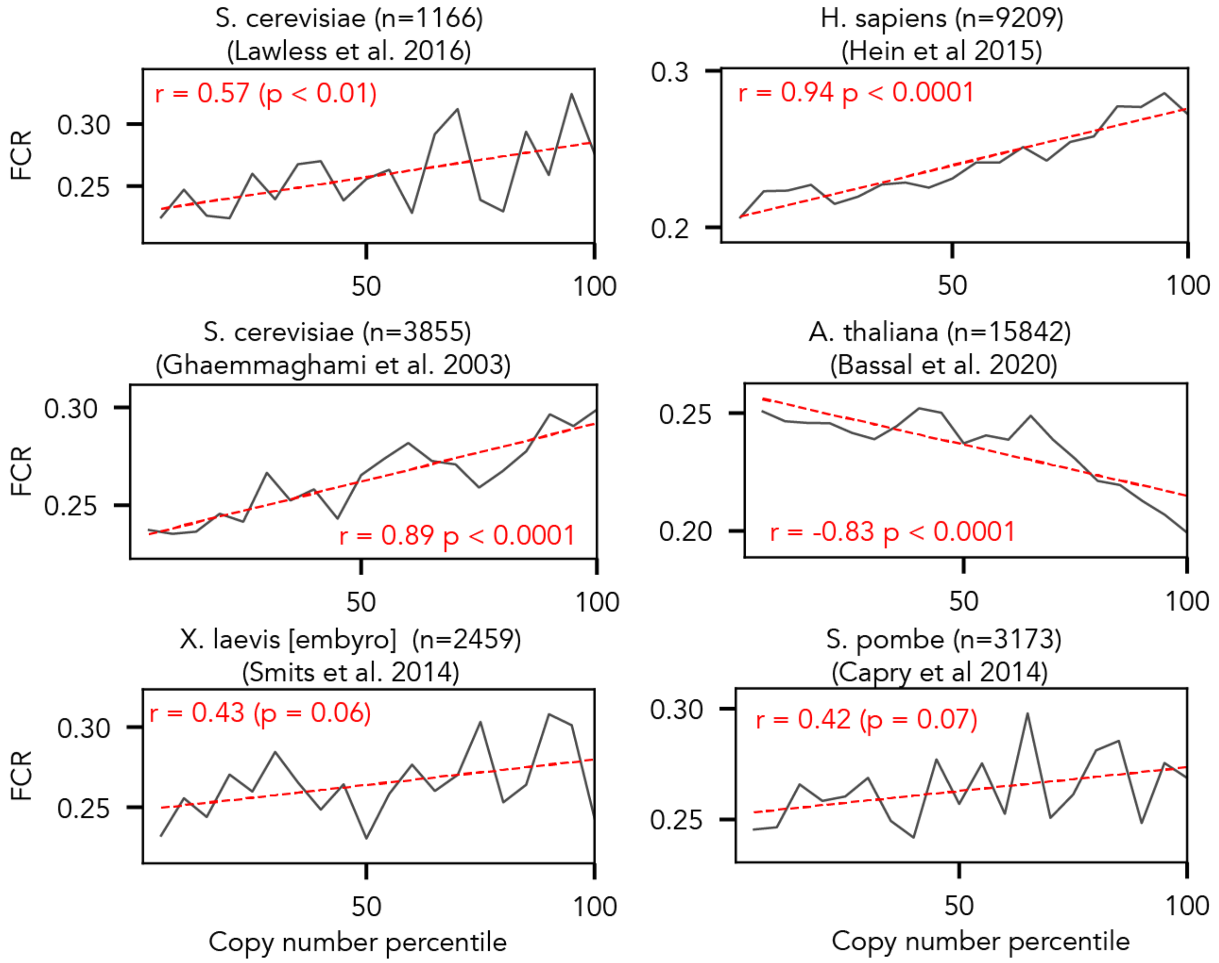


**Figure S5. Copy number vs. the mean fraction of charged residues (FCR) in disordered regions from eight different mass spectrometry datasets**. Taking supplementary information from eight different studies, we correlated the average FCR for disordered regions within each ventile against the copy number ventiles (represented as percentiles for convenience), with correlations calculated using Pearson's correlation coefficient. Intriguingly, while yeast and human proteomes showed a consistent correlation whereby IDRs from highly-expressed proteins are more charged, this trend was not statistically significant in *X. laevis* or S. pombe, and a strong anticorrelation was observed in *A. thaliana*. The *E. coli* datasets had insufficient numbers of IDRs for this analysis to be performable.

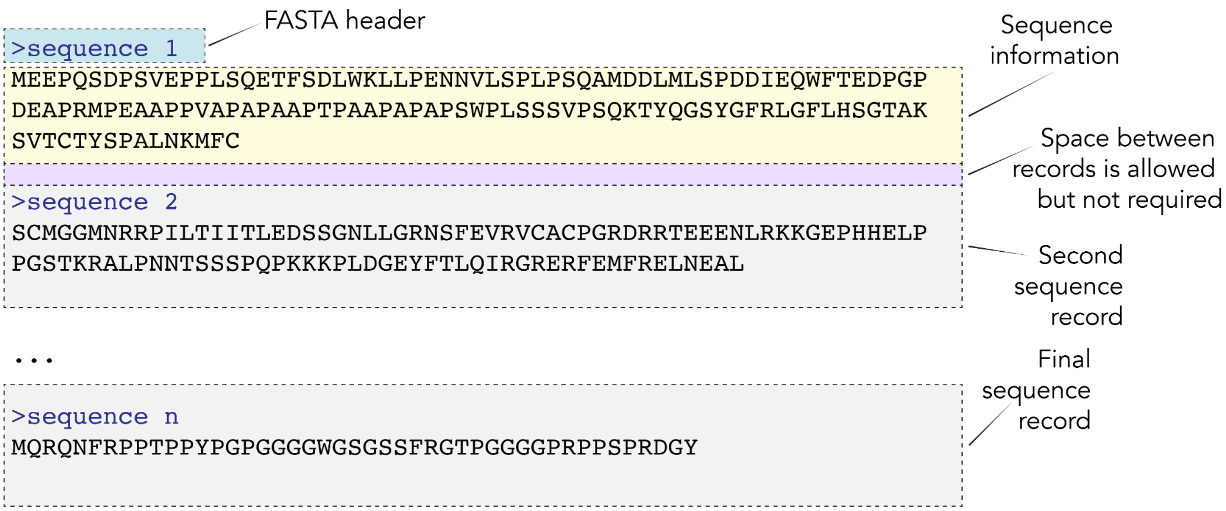


**Figure S6. Schematic of a FASTA file.** FASTA files are a standard format for sharing sequence information. The standard involves a header line (that starts with a “>” symbol) which ideally defines a unique piece of information that reports on the associated sequence. Below the header are one or more lines that contain the protein sequence. The protein record is ended either by the presence of the next header line or by the end of the file.
